# An adaptable, monobody-based biosensor scaffold with FRET output

**DOI:** 10.1101/2022.08.26.505460

**Authors:** Maria F. Presti, Jeung-Hoi Ha, Stewart N. Loh

## Abstract

Protein-based fluorescent biosensors are powerful tools for analyte recognition *in vitro* and in cells. Numerous proteinaceous binding scaffolds have been developed that recognize ligands with affinity and specificity comparable to those of conventional antibodies, but are smaller, readily overexpressed, and more amenable to engineering. Like antibodies, these binding domains are useful as recognition modules in protein switches and biosensors, but they are not capable of reporting on the binding event by themselves. Here, we engineer a small binding scaffold—a consensus-designed fibronectin 3 monobody—such that it undergoes a conformational change upon ligand binding. This change is detected by Förster resonance energy transfer using chemical dyes or cyan and yellow fluorescent proteins as donor/acceptor groups. By grafting substrate recognition residues from different monobodies onto this scaffold, we create fluorescent biosensors for c-Abl Src homology 2 (SH2) domain, WD40-repeat protein 5 (WDR5), small ubiquitin-like modifier-1 (SUMO), and h-Ras. The biosensors bind their cognate ligands reversibly, with affinities consistent with those of the parent monobodies, and with half times of seconds to minutes. This design serves as generalizable platform for creating a genetically-encoded, ratiometric biosensors by swapping binding residues from known monobodies, with minimal modification.

---

Protein-based sensors are used in a wide range of applications, from detecting spatial distributions of molecules and their time-dependent changes in concentration, to identifying disease biomarkers *in vitro* and *in vivo*^1,2^. Biosensors consisting of a single protein molecule typically work by coupling an input signal (ligand binding) to a measurable output by means of a conformational change. The output is often a fluorescence change (ratiometric or intensiometric) since fluorescent proteins (FPs) can easily be incorporated into the design. Proteins are capable of many types of conformational change, including the folding and unfolding reactions featured in this study, and several of these have been used to effect communication between input and output domains.

A major challenge in protein biosensor design is to develop a scaffold that can be easily modified to recognize new molecules of interest. Existing sensors are often exquisitely sensitive and precise, but the mechanisms by which they achieve this response are almost always specific to their intended target and are not readily transferrable to other ligands. Perhaps the pre-eminent example is the GCaMP family of calcium sensors, which consist of calmodulin and one of its binding peptides fused to an FP. By virtue of their strong intensiometric fluorescence change, rapid response times, and genetic encodability, GCaMPs have transformed the study of calcium signaling in cells. Yet this turn-on mechanism has proven difficult to apply to the detection of other targets. GCaMPs take advantage of calmodulin’s peculiar Ca^2+^-induced peptide-binding activity, and while GCaMP-like sensors with recognition domains other than calmodulin have been developed^3–5^, each design represents a new protein engineering effort with substantial screening and optimization required.

The goal of this study is to create a self-reporting protein biosensor that one can customize to bind a new target with minimal modification. Existing examples can be grouped into two categories: those consisting of a single protein and those composed of two or more components. In the latter class, the components come together in the presence of ligand and report on the interaction by either reconstituting a split FP or enzyme (e.g., luciferase or β-lactamase), or bringing Förster resonance energy transfer (FRET) pairs into proximity. The advantage is that, when the system is tuned properly, signal-to-noise ratio tends to be high due to low background levels of enzymatic activity or FRET. The limitations are that it can be challenging to tune the system such that the proteins only assemble in the presence of ligand, and oftentimes the components may need to be present at approximately equal concentrations. Single-molecule sensors remove those limitations, but they generally require the molecule to undergo a binding-induced conformational change, which is relatively rare among naturally-occurring proteins.

One way to introduce allostery into proteins that do not undergo intrinsic conformational change is with the alternate frame folding (AFF) mechanism^6–11^. The AFF mechanism, as well as how to create a functional AFF switch, has been described in detail^12^. Briefly, the full-length biosensor construct consists of the native sequence and a segment duplicated from either terminus, which is fused to the opposite terminus using a peptide linker long enough to span the termini of the parent protein. This procedure is illustrated in Figure 1B using the 93-amino acid fibronectin type III protein (FN3 or monobody; Figure 1A). Duplication creates two mutually-exclusive folding frames (Figure 1B), generating the native fold (N-fold) and circularly permuted fold (cp-fold) (Figure 1C). Binding site mutations are introduced into one of the folds to create exclusivity of binding, while the shared sequence between the two folds ensures exclusivity of folding. Fluorophores are placed in the surface loop at which the native protein was permuted and at either the amino or carboxy terminus, depending on whether the duplicated sequence was appended to the amino or carboxy terminus, respectively. At these positions, the fluorophores are at the amino and carboxy termini of the cp-fold. These positions are proximal in all circularly permuted proteins, and FRET efficiency is consequently high when the switch adopts the cp-fold (Figure 1C). When the sensor switches to the N-fold, the fluorophores become separated by the duplicated segment and FRET efficiency is expected to decrease. The AFF mechanism has been used to create Ca^2+^and ribosebinding biosensors, using calbindin^6^ and RBP^9^ scaffolds respectively, as well as an artificial HIV-1 protease activated zymogen^10^ and a protease-activated, color-changing fluorescent protein^13^.

**Figure 1.**
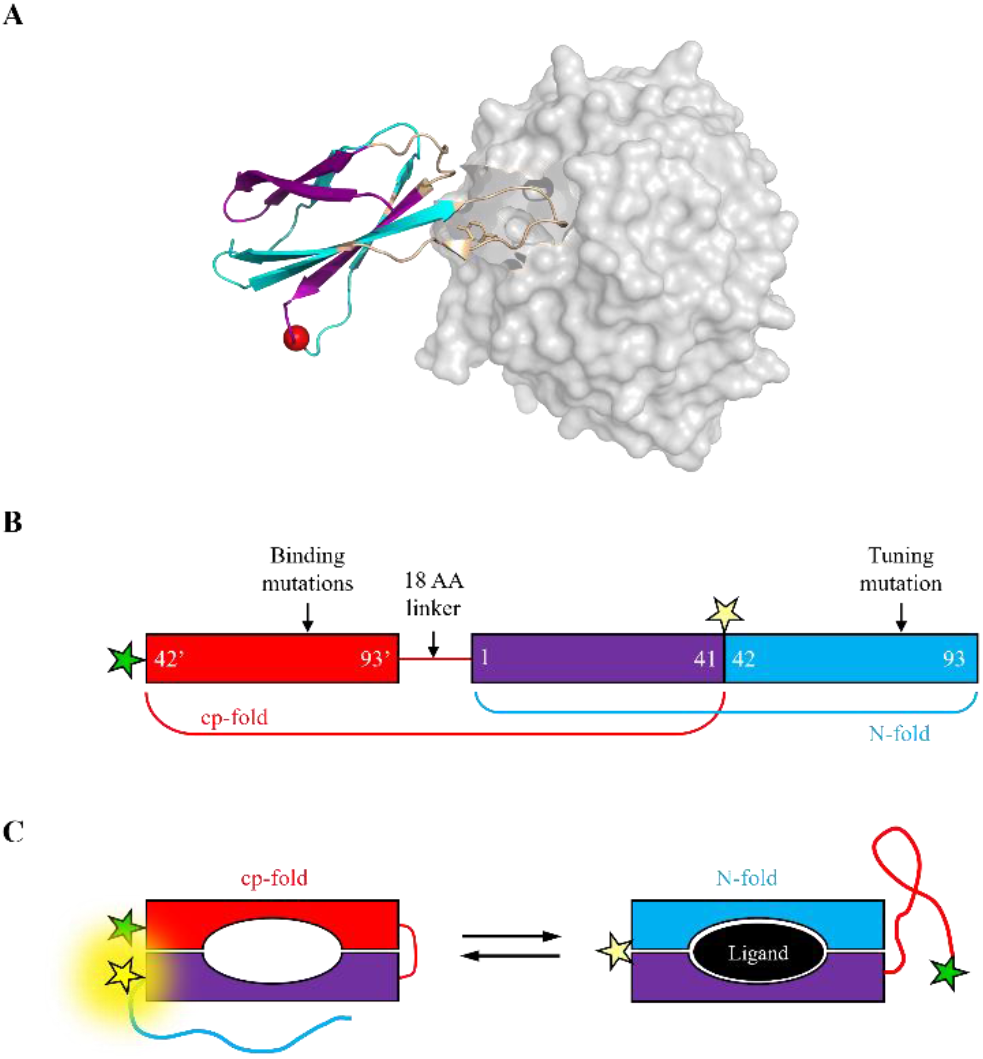
Design of FN3-AFF sensors. **(A)** The structure of FN3 (ribbons) complexed with WDR5 (grey surface) shows the binding interface (BC and FG loops; wheat color), the site of circular permutation (position 42; red sphere), and regions that were duplicated (cyan) and shared (purple) in the FN3-AFF sensor. PDB ID 6BYN. **(B)** FN3-AFF switches were created by copying residues 42 – 93 and appending the duplicate segment (denoted 42’ – 93’) to the N-terminus of FN3, joined by an 18AA linker. The thermodynamic stabilities of the cp-fold (red and purple) and the N-fold (purple and blue) were balanced by introducing a tuning mutation into the N-fold, and ligand binding of the cp-fold was knocked out by introducing binding-null mutations into the red sequence. **(C)** FN3-AFF interconverts between the cp-fold and the N-fold, with the chief difference in their structures being the extension of a C-terminal tail (residues 42 – 93) from the cp-fold and an N-terminal tail (residues 42’ – 93’) from the N-fold. Binding of the ligand (black oval) shifts the population from the cp-fold to the N-fold, resulting in a loss of FRET efficiency as the donor fluorophore (green star) and acceptor fluorophore (yellow star) move apart. FN3-AFF is shown; FN3-AFF^rev^ consists of the binding mutations transferred from the cp-frame to the N-frame.

Here, we create an adaptable and genetically encoded switch by applying the AFF mechanism to the 10^th^ domain of FN3, termed monobody, a small protein that has been evolved by *in vitro* selection methods to specifically bind ∼30 protein targets^14,15^ including epidermal growth factor receptor^16^, maltose-binding protein (MBP)^17^, mixed lineage kinase domain-like protein ^18,19^, and severe acute respiratory syndrome coronavirus 2 spike protein^20,21^. Like the antibody light chains they resemble, monobodies achieve molecular recognition by presenting different residues at surface positions (1-3 CDR-like loops as well as several βstrands) while maintaining a constant amino acid sequence at other positions^14^. Monobodies have been developed for use in degradation systems^22–25^, light-controlled applications^26,27^, cancer therapeutics^28–30^, and as inhibitors^14,31–33^ and biosensors^34,35^.

Monobodies are excellent binders, but they are not allosteric switches, and thus are not able to self-report binding events. We convert FN3 to a switchable protein scaffold using the aforementioned AFF mechanism, and, using crystal structures of known monobody-target complexes, generate fluorescent biosensors that recognize four human protein targets: cAbl Src homology 2 domain (SH2)^31,36^, small ubiquitin-like modifier-1 (SUMO)^36^, WD40-repeat protein 5 (WDR5)^33^, and h-Ras^37^. FRET output is established by chemical labeling with BODIPY fluorescence donor and acceptor groups, or by fusion of cyan and yellow fluorescent proteins to create a fully genetically-encoded sensor with ratiometric response.

## Results

### Generation of monobody binding scaffolds

The structure of FN3 consists of a seven-stranded beta sandwich (strands A – G) with the BC, DE, and FG loops corresponding to antibody CDR regions (Figure 1A). Because circularly permuting a protein is often destabilizing, we based our switch on FN3 that had been previously stabilized using the consensus design method^38^. We first converted this naïve, consensus-designed FN3 (PDB 4U3H) into an SH2-recognizing monobody by examining the crystal structure of the original (non-consensus designed) HA4 monobody in complex with SH2 (PDB 3K2M) and identifying which amino acids on HA4 were in contact with SH2. In addition to the three CDR-like loops, four amino acids in strand F of HA4 interact with SH2. We then transferred these four residues and grafted the entire BC, DE, and FG loops from HA4 onto the consensus-designed FN3. This SH2-recognizing, consensus-designed FN3 sequence served as the template for creating WDR5 and SUMO monobodies by the same process (see Methods). The resulting monobodies are designated with superscripts indicating their respective target ligands (FN3^SH2^, FN3^WDR5^, and FN3^SUMO^) (Table 1). Finally, for labeling with fluorescent dyes, a Cys residue was introduced at the position in the CD loop at which FN3 was circularly permuted (*vide infra*). For simplicity, we designate this as position 42 for all sensors, although the numbering shifts by several residues depending on the FN3 construct. Amino acid sequences of all constructs created for this study are shown in Supplemental Figure S1.

To create circular permutants, we chose to make the new amino terminus position 42 (red sphere in Figure 1A) because the CD loop is not varied in most directed evolution procedures, and previous studies have shown that this loop can tolerate cleavage and mutation without undue loss of stability or function^34,39^. We then linked the last residue of the original sequence to the first residue of the original sequence by means of an 18-AA flexible linker long enough to span the ∼36 Å carboxy-to-amino terminal distance of FN3. The sequence terminated with residue 41 and a Cys residue was placed at the amino terminus for labeling purposes. The circular permutants are designated cpFN3 (Table 1).

**Table 1.** Nomenclature of constructs used in this study.

| Name | Description |
| --- | --- |
| FN3 | Monobody |
| cpFN3 | Circular permutant of monobody |
| FN3<sup>bn</sup>, cpFN3<sup>bn</sup> | FN3 and cpFN3 with binding-null mutations introduced |
| N-fold | Native fold of FN3-AFF biosensor (purple and blue regions of Figure 1A) |
| cp-fold | Circularly permuted fold of FN3-AFF biosensor (red and purple regions of Figure 1A) |
| FN3-AFF | Biosensor with binding-null mutations in the cp-fold |
| FN3-AFF<sup>rev</sup> | Biosensor with binding-null mutations in the N-fold |
| FN3-AFF<sup>GE</sup> | Genetically-encoded FN3-AFF<sup>WDR5</sup>, with Cy-Pet and cpYFP fused at positions indicated in Figure 1. |

### Thermodynamic tuning of sensor components

Prior to assembling the AFF sensor, the thermodynamic stabilities (ΔG_fold_) of FN3 and cpFN3 must be approximately balanced. If the binding-competent fold is much more stable than the binding-null fold, then the switch will always be ON regardless of ligand. If the situation is reversed, then high concentrations of ligand will be required to push the equilibrium from OFF to ON states, reducing the sensitivity of the sensor. Guanidine hydrochloride (GdnHCl) denaturation experiments revealed FN3 was much more stable than cpFN3 (Supplemental Figure S2C). Introducing a cavity-creating mutation into a buried position in the unshared region of FN3 (Phe→Gly at position 69 in FN3^SUMO^ and FN3^WDR5^, and at the analogous position 71 in FN3^SH2^) resulted in ΔG_fold_ values of 6.97 ± 0.13 kcal mol^-1^ (FN3^WDR5^), 7.22 ± 0.13 kcal mol^-1^ (FN3^SH2^), and 6.01 ± 2.19 kcal mol^-1^ (FN3^SUMO^) (Supplemental Figure S2 and Supplemental Table 1). These values were approximately equal to those of the corresponding cpFN3 constructs (without the Phe→Gly mutation and containing binding-null mutations; Supplemental Figure S2 and Supplemental Table 1). Thus, no further stability tuning was necessary.

### Functional characterization of sensor components

We determined ligand binding affinities by titrating in ligand and monitoring the increase in fluorescence anisotropy (FA) of fluorescein-labeled FN3 and cpFN3 (Supporting Figure S3). All FN3 and cpFN3 constructs bound their ligands with sub-μM K_d_ (Table 2). K_d_ values of FN3^SH2^ and FN3^WDR5^ were higher than those reported for the parental monobodies by 8-fold and 68-fold, respectively, whereas FN3^SUMO^ bound SUMO with approximately the same affinity as the parental monobody (Supplemental Figure S2A). Circular permutation did not appreciably affect binding affinities in the case of SH2 and WDR5 (Supplemental Figure S2B; Table 2), but K_d_ of cpFN3^SUMO^ was ∼12-fold higher than that of FN3^SUMO^.

**Table 2.** Binding affinities of the FN3 constructs and FN3-AFF switches.

| Construct | SH2<br>Kd ($\mu\text{M}$) | SUMO<br>Kd ($\mu\text{M}$) | WDR5<br>Kd ($\mu\text{M}$) |
| --- | --- | --- | --- |
| FN3 | $0.18 \pm 0.75$ | $0.05 \pm 0.02$ | $0.34 \pm 0.03$ |
| cpFN3 | $0.22 \pm 0.08$ | $0.62 \pm 0.20$ | $0.34 \pm 0.03$ |
| FN3-AFF | $1.05 \pm 0.61$ | $1.92 \pm 0.16$ | $1.38 \pm 0.58$ |
| FN3-AFF<sup>rev</sup> | $3.60 \pm 2.05$ | $1.52 \pm 0.96$ | $0.26 \pm 0.08$ |
| Parental | $0.022^{31}$ | $0.068^{32}$ | $0.005^{33}$ |

For the AFF switch design, it is necessary to knock out function of either the Nfold or the cp-fold. In monobodies, the FG loop interacts extensively with all the ligands and typically contains at least one essential binding residue^34^. We introduced the binding-null mutations Y83A+M84A (SH2), H74G+W76G (SUMO), and R79A (WDR5) into FN3, as well as into cpFN3 at the identical positions, to create binding-null mutants (designated FN3^bn^ and cpFN3^bn^). FA experiments confirmed that FN3^bn^ and cpFN3^bn^ did not interact with their respective ligands (Supplemental Figure S3).

### FN3-AFF sensor performance characterized by BODIPY fluorescence

FN3-AFF^SH2^, FN3-AFF^SUMO^, and FN3-AFF^WDR5^ biosensors were constructed as described in Figure 1A. The binding-null mutations were introduced into the cp-fold, and cysteines from FN3 (internal position 42) and cpFN3 (amino terminus) were carried over to FN3-AFF for labeling purposes. Sensors were purified from *E. coli* and were >95 % pure and monomeric as determined by size exclusion chromatography (SEC) (Supplemental Figure S4A). FN3-AFF biosensors were labeled using BODIPY-FL maleimide. In the cp-fold, the BODIPY groups are at effectively adjacent positions and FRET efficiency (manifested by quenching) is high. In the N-fold, the dyes become separated (Figure 1C) and FRET efficiency is expected to decrease, leading to dequenching. FN3-AFF^SH2^, FN3-AFF^SUMO^, and FN3-AFF^WDR5^ sensors all exhibited an increase in BODIPY emission upon addition of ligand, consistent with the expected cp-to-N fold shift (Figure 2A). The three sensors bound their ligands with K_d_ values 4 – 40-fold higher than those of the respective, isolated FN3 proteins. This trend is due at least in part to a portion of the binding energy necessarily being used to drive the cp-to-N conformational change^6–8,10^.

The observed increase in BODIPY fluorescence is consistent with the AFF-mediated conformational change, but it’s possible that it could be due to binding-induced dequenching of BODIPY in the absence of a fold shift. To distinguish between these scenarios, we reversed the switch direction by transferring the binding-null mutations from the cp-fold to the N-fold, thereby creating N-to-cp-fold switches (FN3-AFF^rev^). Fluorescence binding curves of FN3AFF^rev^ sensors appeared as mirror images of their FN3-AFF counterparts (Figure 2B), indicating that binding induced a shift from low-FRET (N-fold) to high-FRET (cp-fold) states. This finding confirms the conformational change had indeed reversed and was caused by the AFF-mediated fold shift. Reversing the switch direction did not alter the binding affinities of SH2 and SUMO, but it increased affinity of WDR5 by ∼5-fold (Figure 2 and Table 2). This latter result suggests that the binding-null mutations slightly destabilized the N-fold in FN3-AFF^WDR5,rev^, increasing the initial population of the cp-fold and reducing the binding energy required to bring about the N-to-cp fold shift.

**Figure 2:**
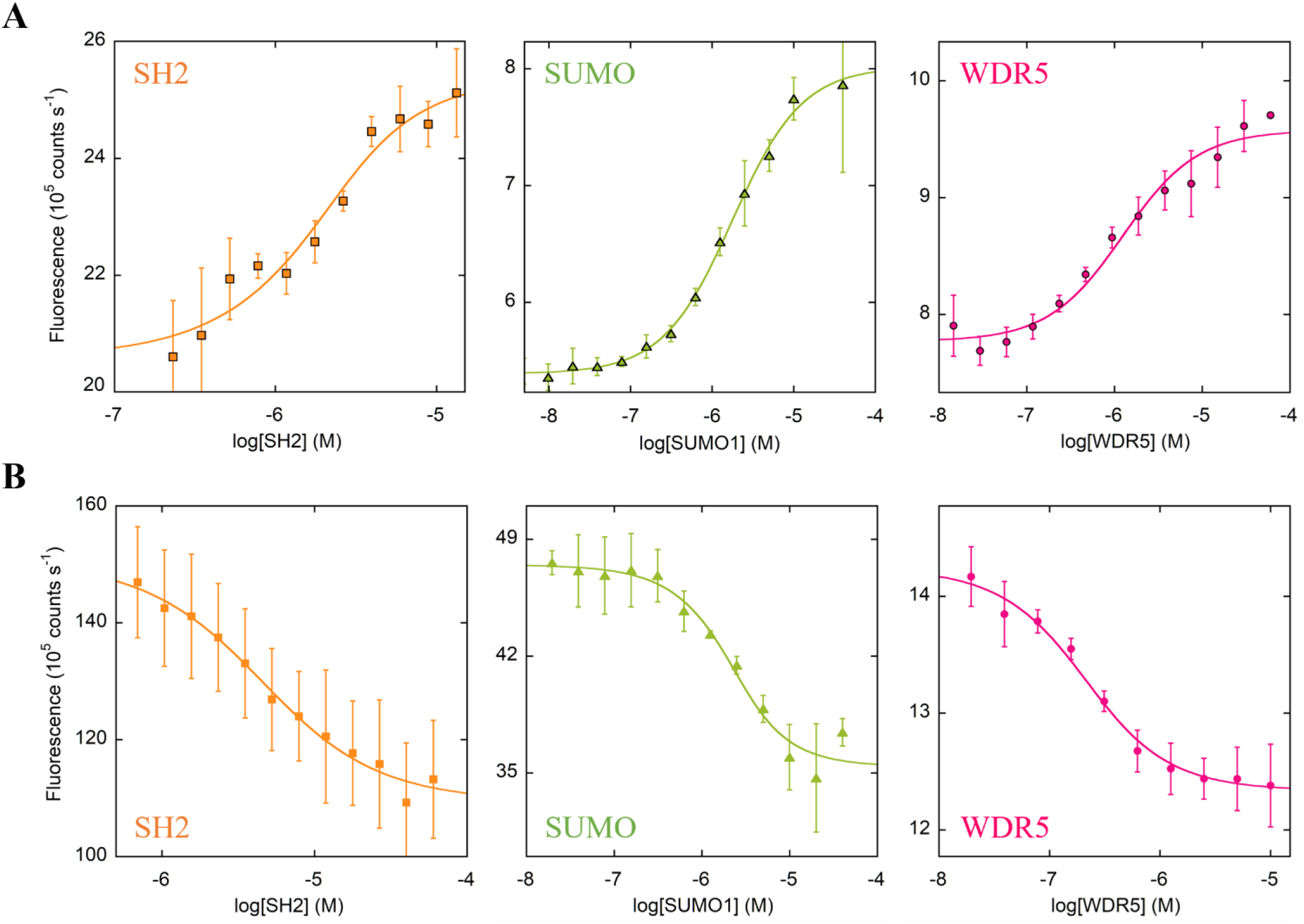
Equilibrium binding of FN3-AFF sensors (A) and FN3-AFFrev sensors (B) characterized by BODIPY FRET. Fluorescence intensity was monitored at 510 nm. Solid lines indicate best fits to the one-site binding equation. Errors are s.d. (n = 3).

### Rapid generation of a new FN3-AFF sensor for h-Ras

Having developed biosensors for detecting SH2, SUMO, and WDR5, we asked whether a new sensor could be made by changing the ligand binding residues of the existing FN3AFF scaffold without additional thermodynamic balancing or other optimization. We chose to target h-Ras for this test. Ras is frequently mutated in cancer and Ras-targeting sensors could potentially aid in diagnosing cancers with mutant Ras and screening Ras inhibitors in cells. We looked at the crystal structure of an h-Ras-targeting monobody (PDB 5E95), termed NS1, and grafted the binding residues from the FG and BC loops as well as three residues in strand D onto our existing FN3-AFF^SH2^ sensor (Supplemental Figure 1). Addition of h-Ras produced an increase in BODIPY fluorescence, consistent with the expected conformational change, with K_d_ = 423 ± 200 nM (∼26-fold weaker than NS1)^37^ (Supplemental Figure S5). This result suggests that it should be possible to create additional sensors by following the same procedure.

### Construction and characterization of a genetically-encoded WDR5-binding FN3-AFF sensor

To create the genetically-encoded FN3-AFF^WDR5^ biosensor (designated FN3-AFF^GE^), we replaced the amino terminal Cys with a cyan fluorescent protein (CyPet)^42^ and the Cys at internal position 42 of the N-fold with a circularly-permuted yellow fluorescent protein (cpYFP) (Supplemental Figure S1B). Circular permutation was necessary to reduce the amino-to-carboxy terminal distance of YFP, so that it could be inserted into the surface loop of FN3 without unfolding FN3 and thus induce mutually exclusive folding of the YFP and FN3 domains of the fusion protein^40,41^. cpYFP was generated by permuting Clover fluorescent protein at position 195^43^ and introducing F46L, F64L, V67L, and H203Y mutations^44^ to change its fluorescence from green to yellow as described in our earlier work^45^. To determine whether inserting cpYFP into FN3^WDR5^ affected the function of either domain, we purified the FN3^WDR5^-cpYFP fusion protein and determined it was brightly fluorescent and bound WDR5 with K_d_ = 727 ± 53 nM (Supplemental Figure S6). As with the other sensors, FN3-AFF^GE^ expressed in *E. coli* as a soluble protein and was monomeric by SEC (Supplemental Figure S4B).

Upon adding WDR5, fluorescence spectra of FN3-AFF^GE^ revealed the ratiometric increase in CyPet emission and decrease in cpYFP emission expected for the high-FRET to low-FRET change of the cp-to-N fold shift (Figure 3A). Fitting the acceptor fluorescence (525 nm) to the one-site binding equation yielded K_d_ = 723 ± 120 nM and fitting acceptor/donor fluorescence (525 nm/475nm) yielded a 1.31-fold decrease in FRET ratio (Figure 3C). To validate that the observed FRET change was due to the AFF-mediate fold shift, we transferred the binding mutations from the cp-fold to the N-fold (FN3-AFF^GE,Rev^) and observed the expected ratiometric decrease in CyPet emission and increase in cpYFP emission upon addition of WDR5 (Figure 3B). Fitting the acceptor fluorescence gave K_d_ = 351 ± 64 nM, and FRET efficiency, as measured by acceptor/donor fluorescence, increased by 1.17-fold (Figure 3C).

**Figure 3.**
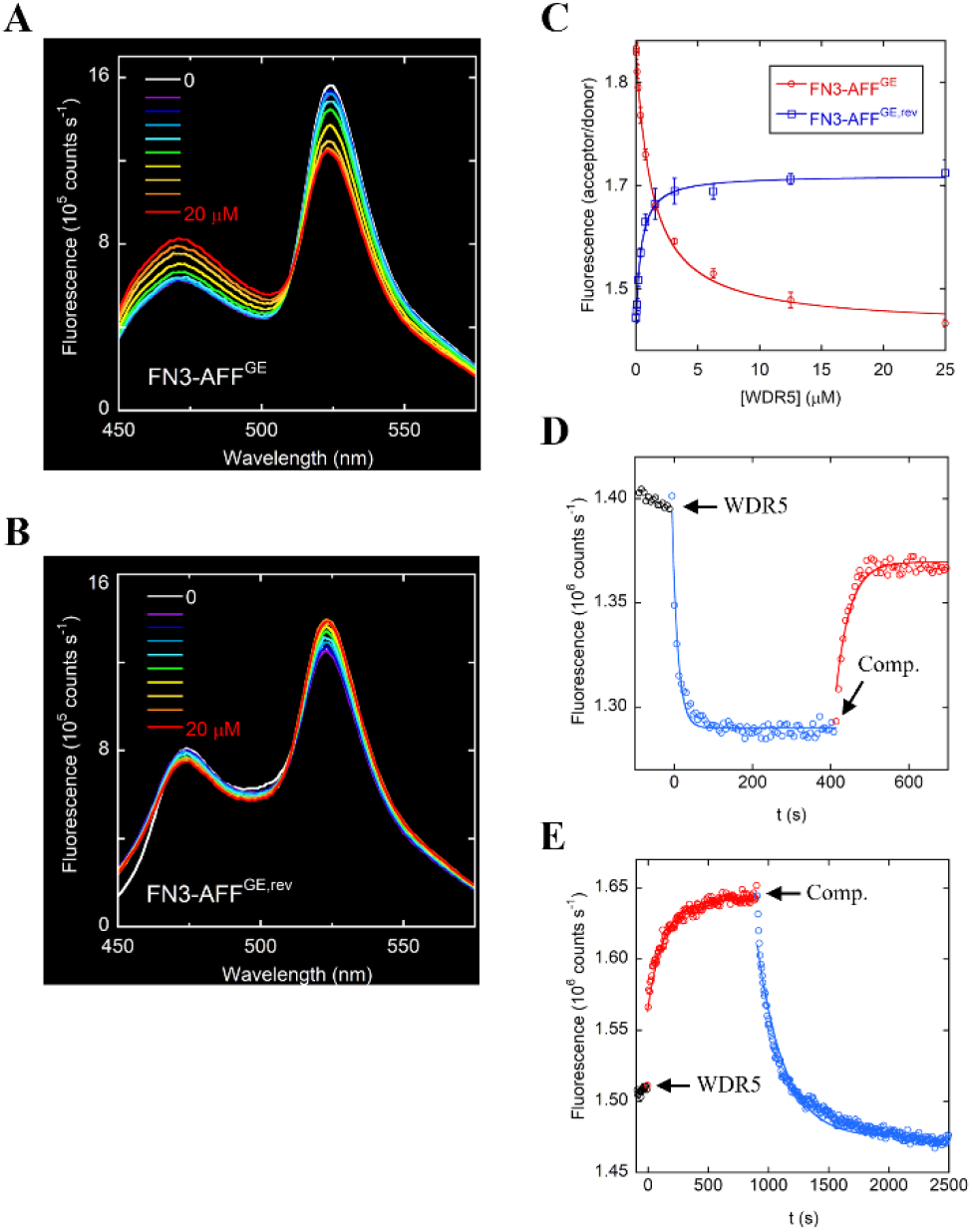
FRET response of FN3-AFF^GE^ and FN3-AFF^GE,rev^. Raw fluorescence spectra of FN3-AFF^GE^ **(A)** and FN3-AFF^GE,rev^ show the expected decrease and increase in FRET efficiency, respectively, with addition of WDR5. Concentrations of WDR5 from 0 to 20 μM (increasing by 2-fold increments) are indicated in the insets. Lines are best fits to the one-site binding equation and error bars are s.d. (n = 3). **(C)** FN3-AFF^GE^ (red) and FN3-AFF^GE,rev^ (blue) bind WDR5 with sub-μM affinity. FN3AFF^GE^ **(D)** and FN3-AFF^GE,rev^ **(E)** bind WDR5 rapidly and reversibly. Acceptor fluorescence was monitored after adding 10 µM WDR5 and 50 µM competitor (FN3^WDR5^) at the times indicated by arrows. Blue and red data represent cp-to-N and N-to-cp fold shifts, respectively. Lines are best fits to single-exponential equations.

The turn-on rate (k_ON_) of FN3-AFF^GE^ was obtained by mixing the sensor with WDR5 and fitting the change in FRET efficiency as reported by the decrease in cpYFP fluorescence. The data fit well to a single exponential function, yielding k_ON_ = 12.5 ± 0.9 min^-1^ (Supplemental Figure S7). The k_ON_ rate did not depend on WDR5 concentration over the range of 5 – 20 μM. This result, together with the apparent one-step binding reaction, suggests a conformational capture mechanism in which the rate limiting step was most likely the cp-to-N conformational change, with WDR5 binding rapidly to the N-fold.

To investigate the reversibility of FN3-AFF^GE^, and to measure the turn-off rate (k_OFF_), we repeated the binding experiment and then added excess competitor (FN3^WDR5^) to the FN3-AFF^GE^/WDR5 complex. Reversibility was evaluated by the increase in cpYFP fluorescence intensity. Fluorescence returned to close to the original value with k_OFF_ = 1.56 ± 0.18 min^-1^ (Figure 3D), demonstrating that the switch is reversible. Finally, we measured k_ON_ and k_OFF_ of FN3AFF^GE,rev^ by the same method. As anticipated, the directions of the observed FRET intensity changes were opposite to those of FN3-AFF^GE^ (Figure 3E), confirming the AFF-mediated conformational change and demonstrating that FN3-AFF^GE,rev^ also switched reversibly. k_ON_ (0.24 ± 0.12 min^-1^) and k_OFF_ (0.3 ± 0.03 min^-1^) were slower than their respective values for FN3-AFF^GE^.

## Discussion

The adaptability of our sensor is enabled by basing the design on an established, versatile binding domain. Several other groups have applied the same principle to create protein switches that can be made to recognize other targets. Although they are not biosensors, opto-monobodies^26^ and moonbodies^27^ use light to control the binding affinity of monobodies. The *Avena sativa* LOV2 domain was inserted into the DE (OptoMb) or EF (moonbody) loops of the SH2binding HA4 monobody^36^. Absorption of blue light induces the Jα helix of LOV2 to unfold, which resulted in loss of SH2 binding affinity. Opto-nanobodies^46^ and sunbodies^27^ were created using the same LOV2 insertion strategy applied to nanobodies. Of similar size and structure to monobodies, nanobodies are single-chain camelid antibodies that can be evolved to bind custom targets by standard hybridoma technology. Depending on the nanobody loop into which LOV2 was fused, light caused ligand binding affinity to decrease or increase^46^. This result underscores the lack of mechanistic understanding of the switching mechanisms and the need for screening different insertion sites and LOV2 truncation mutants. In addition to SH2, photoactivatable binders for MBP, eGFP, SUMO, mCherry, and F-actin were generated by inserting LOV2 into the respective monobodies or nanobodies.

Using an *E. coli* library screening method based on antibiotic resistance, Ostermeier and colleagues approached developing an adaptable biosensor by inserting an enzymatic output domain, circularly permuted TEM-1 β-lactamase (cpBLA), into random locations in FN3 and a designed ankyrin repeat protein (DARPin), both of which had been previously evolved to bind MBP^47^. DARPins are another class of small antibody mimetics whose structure consists of tandem repeats of paired α-helices connected by loops that comprise the ligand binding interface^48,49^. BLA enzymatic activity of both switches increased ∼10-fold in the presence of MBP, although binding affinity of MBP was substantially diminished compared to the parental monobody and DARPin. The authors then mutated the monobody domain to recognize eGFP and ySUMO and the DARPin domain to bind eGFP and APH(3’)IIIa. The monobody-based switches failed to show the expected changes in specificity, as did the eGFP DARPin sensor, but the DARPin APH(3’)IIIa sensor exhibited a 25-fold increase in activity in the presence of APH(3’)IIIa and no change when MBP was added. One distinction between our study and that of the Ostermeier group is that the allosteric mechanisms by which the latter switches operate are unknown. Monobodies and DARPins do not undergo appreciable ligand-induced conformational changes, so it was unclear how the binding signal is transmitted to the cpBLA domain. This point illustrates how an efficient selection tool—one conferring antibiotic resistance in this case—can greatly facilitate bioswitch discovery.

Baker and coworkers recently introduced an adaptable two-component biosensor called lucCage^50^ that was based on their earlier LOCKR design^51^. Instead of achieving molecular recognition via a naturally-occurring binding domain, lucCage used a de novo designed six-helix bundle composed of the N-terminal, five-helix ‘cage’ and the C-terminal, single-helix ‘latch’. Embedded into the latch was a short fragment (smallBit) from the luminescent protein nanoluciferase^52^. Using a combination of computational and experimental screening methods, they modified the C-terminus of the latch such that it recognized multiple target ligands (BCL-2, IgG1 Fc domain, HER2 receptor, and several others). The exogenously-added ‘key’ consisted of a second copy of the latch (without smallBit or the ligand binding sequence) fused to largeBit, the nanoluciferase fragment complementary to smallBit. Binding of the target weakened the interaction between latch and cage, causing the latch to exchange with the key. Complementation of largeBit with the newly-exposed smallBit established luminescent output. To achieve switching functionality, the thermodynamic stability of the system was tuned by mutation so that the latch was poised to dissociate from the cage but did not do so until the target bound, thus minimizing false activation while maximizing target affinity. LucCage was the first biosensor to be designed in silico, and whose binding specificity was altered by computational redesign.

The lucCage mechanism is analogous to that of the earlier fragment exchange (FREX) biosensor based on the FN3 scaffold^34^. The FREX sensor consisted of eGFP fused to the N-terminus of the HA4 monobody, which harbored the same binding-null mutations in our current work plus a thermodynamic tuning mutation. The tuning mutation, which was either Val, Ala, or Gly in place of the hydrophobic core residue Ile75, served to progressively destabilize FN3 without unfolding it, just as in the lucCage tuning process. The ‘key’ in this case was an exogenously-added peptide consisting of residues 48 to 100 of HA4, which contained the WT sequence at the binding and tuning sites and was fused to mCherry. Binding of SH2, in combination with restoration of Ile at position 75, caused the exogenous fragment to exchange with the endogenous fragment, producing a robust FRET increase in cells and in vitro. FREX is thus the bimolecular version of FN3-AFF. Because the fluorophores were on separate molecules that came together only in the presence of target, background FRET signal was minimal and the ratiometric change was unusually strong (up to 8.6-fold). However, approximately equal concentrations of donor and acceptor fluorophore are required for maximal FRET response, and while it is straightforward to maintain this condition in vitro, it may be difficult to do so in cells.

The attributes of the FN3-AFF sensor are that it is unimolecular, genetically encoded, ratiometric, reasonably fast, reversible, and customizable. It should be possible to rationally mutate FN3-AFF^GE^ to bind new targets, provided that a high-resolution structure exists of the monobody-ligand complex. Although mutations are generally limited to surface-exposed residues, this procedure can still perturb the thermodynamic balance between the N-fold and cp-fold. If so, an additional tuning mutation at Ile75 or Ile75’ as described in the FREX study can be introduced to restore balance.

There are two main limitations of the current FN3-AFF design. First, ligand affinity is typically lower than that of the parental monobody. Some binding energy is inevitably siphoned off to drive the fold shift. The main reason, however is that transposing the ligand-contacting residues from different monobodies onto a common scaffold does not capture all the nuances of the binding interactions^38,53^. This method can reproduce parental binding affinities (e.g., FN3^SUMO^ (Table 2) and a VEGFR2-binding consensus-designed FN3^53^), but these cases seem to be exceptions. Additional computational redesign, such as that employed in the lucCage study, may be expected to improve affinity. Alternately, it may be more facile to create new FN3-AFF sensors by applying the protein engineering steps outlined in Figure 1 to pre-existing monobodies, as was done in the optoMB and moonbody studies. Sites of binding-null mutations, tuning mutation, and circular permutation can be deduced by sequence alignment in the absence of a known structure, and the same 18-AA linker can be used.

The second limitation of FN3-AFF is one experienced by most small, unimolecular FRET-based sensors: the donor-acceptor distance usually does not change by a large amount between free and bound states. It may be possible to improve the FRET response of our sensor by introducing peptide linkers between one or both fluorescent groups and the protein, to extend the donor-acceptor distance in the Nfold.

In conclusion, we created an adaptable biosensor by engineering allostery in a consensus-designed monobody scaffold using the alternate frame fold mechanism. By balancing the stabilities of the Nand cp-folds and modifying the CDR-like loop and beta strands of FN3, we made FN3AFF biosensors for SH2, WDR5, and SUMO. A fourth sensor for h-Ras was generated by mutating the binding interface only, without additional optimization. The fold switch occurred within seconds to minutes and was fully reversible. The molecules and methodology herein provide a template for non-experts to generate additional genetically-encoded biosensors for targets of choice.

## Experimental Procedures

### Generation of WDR5, SUMO, and h-Ras binding monobodies

FN3^SUMO^ was made by grafting the BC and FG loops from monobody SL8^36^ onto FN3^SH2^. FN3^WDR5^ was created by grafting the BC and FG loops, as well as four residues from strand D, from the Mb(S4) monobody^33^ onto FN3^SH2^. FN3^Ras^ was generated by grafting the BC and FG loops, and four residues strand D, from the NS1 monobody^37^ onto FN3^SH2^. Full amino acid sequences of these and all constructs are listed in Supplemental Figure S1.

### Gene Construction and Protein Purification

The c-Abl SH2 plasmid was a gift from S. Koide (NYU Langone Health, New York, NY). pET11a-SUMO1 was a gift from Frauke Melchior (Addgene plasmid #53138; http://n2t.net/addgene:53138; RRID:Addgene_53138)^54^. WDR5 plasmid was a gift from M. Cosgrove (Upstate Medical University, Syracuse, NY). CyPet gene was kindly provided by Patrick Daugherty (University of California, Santa Barbara). FN3^SH2^ and h-Ras genes were synthesized by GenScript (Piscataway, NJ). cpFN3^SUMO^ and cpFN3^WDR5^ genes were synthesized by Eurofins Genomics (Louisville, KY). Full amino acid sequences of all constructs are listed in Supplemental Figure S1. All genes were cloned into the pET41b *E. coli* expression vector and fully sequenced.

*Escherichia coli* BL21(DE3) cells were transformed with the plasmids described above. Cultures were grown in LB medium at 37 °C to OD_600_ ∼0.6, induced with IPTG, and shaken for an additional 18 – 20 h at 18 °C. FN3-AFF sensors were purified from the soluble fractions of cell lysates using nickel nitriloacetate resin (Gold Biotechnology, St. Louis, MO) following the manufacturer’s protocols. Proteins were then further purified using a Superdex-75 column (Cytiva, Marlborough, MA) equilibrated in 20 mM Tris (pH 7.4), 0.3 M NaCl, 1 mM tris(2-carboxyethyl)phosphine (TCEP), and 0.005% Tween-20. SH2^34^, SUMO^54^, and WDR5^55,56^ were purified as described. h-Ras did not contain a HisTag and was purified on a Q-Sepharose column (Bio-Rad) in 20 mM Tris (pH 7.4), 10 mM NaCl, 5 mM MgCl_2_, 0.1 mM TCEP, 0.005 % Tween-20, using a 0.01 – 1 M NaCl gradient, then passing it through the Superdex-75 column as above with the addition of 5 mM MgSO_4_. All proteins were judged to be >95% pure by SDS-PAGE.

### Chemical Dye Labeling

Proteins were reduced with 1 mM TCEP for 1 h then passed through a 10DG desalting column (Bio-Rad, Hercules, CA) to remove the TCEP. A 2-fold excess of BODIPY-FL C_5_-maleimide (FN3-AFF samples) or Fluorescein 5 maleimide (FN3 and cpFN3 samples) (ThermoFisher Scientific, Waltham MA) was immediately added and the reaction was allowed to proceed for 1 h at room temperature or overnight at 4 °C. Excess dye was removed by desalting as described above. Final protein concentrations and labeling efficiencies of FN3-AFF proteins were calculated using the BODIPY-FL molar absorptivity (80,000 M^-1^cm^-1^ at 504 nm) and absorbance correction factor (0.04 at 280 nm) provided by the manufacturer. BODIPY-FL labeling efficiency of FN3AFF sensors was typically 30 – 45 %.

### Equilibrium Binding Experiments

BODIPY-labeled FN3-AFF (100 nM) was mixed with the ligands at the indicated concentration and equilibrated for 2 – 4 h at 22 °C before being read on a FluoroMax-4 spectrofluorometer (Horiba, Kyoto, Japan). Scan settings were 490 nm excitation and 500 – 600 nm emission. Binding buffer was 20 mM Tris (pH 7.4), 0.3 M NaCl, 0.005% Tween-20, 0.1% BSA, and 1 mM TCEP. BODIPY fluorescence at 510 nm was plotted as a function of [ligand] and fitted to the quadratic binding equation to obtain K_d_. FN3-AFF^GE^ binding experiments were performed as above, except fluorescence spectra were recorded on a SpectraMax i3x plate reader (Molecular Devices, San Jose, CA) with excitation at 435 nm and emission from 460 – 650 nm. K_d_ was determined by fitting acceptor fluorescence only (525 nm)^57^ to the quadratic binding equation.

### FN3-AFF^GE^ Kinetic Experiments

Sensor turn-on and turn-off kinetics were tracked by exciting CyPet (435 nm) and monitoring YFP emission (525 nm) on the spectrofluorometer. FN3-AFF^GE^ (0.5 μM) was mixed with 10 μM WDR5 at time zero, after recording the sensor-only baseline signal for ∼2 min. After binding was complete, 50 μM FN3^WDR5^ competitor was added. Data shown in Figure 3D and Figure 3E were multiplied by the appropriate factors to account for sensor dilution. k_ON_ and k_OFF_ rates were obtained by fitting the binding and unbinding data to single exponential functions.

## ASSOCIATED CONTENT

## Supporting Information

Amino acid sequences of FN3-AFF switches, stability data of N-fold and cp-fold, binding data of Nfold and cp-fold and validation of binding null mutations, FN3-AFF SEC characterization, FN3-AFF^Ras^ binding data, FN3^WDR5^ with inserted cpYFP binding data, concentration dependence kinetics of FN3-AFF^GE^.

## AUTHOR INFORMATION

### Author Contributions

The manuscript was written by M.F.P. and S.N.L. All authors have given approval to the final version of the manuscript.

### Notes

The authors declare no competing financial interest.

## ABBREVIATIONS

FN3: monobody;
AFF: alternate frame folding;
SH2: c-Abl Src homology 2 domain;
WDR5: WD40-repeat protein 5;
SUMO: small ubiquitin-like modifier-1;
GE: genetically-encoded;
N(-fold): native;
cp(-fold): circularly permuted

## REFERENCES

(1) Tamura, T.; Hamachi, I. Recent Progress in Design of Protein-Based Fluorescent Biosensors and Their Cellular Applications. ACS Chem. Biol. 2014, 9 (12), 2708–2717. https://doi.org/10.1021/cb500661v.

(2) Kim, H.; Ju, J.; Lee, H. N.; Chun, H.; Seong, J. Genetically Encoded Biosensors Based on Fluorescent Proteins. Sensors 2021, 21 (3), 795. https://doi.org/10.3390/s21030795.

(3) Nadler, D. C.; Morgan, S.-A.; Flamholz, A.; Kortright, K. E.; Savage, D. F. Rapid Construction of Metabolite Biosensors Using Domain-Insertion Profiling. Nat. Commun. 2016, 7, 12266. https://doi.org/10.1038/ncomms12266.

(4) Ibraheem, A.; Campbell, R. E. Designs and Applications of Fluorescent Protein-Based Biosensors. Curr. Opin. Chem. Biol. 2010, 14 (1), 30–36. https://doi.org/10.1016/j.cbpa.2009.09.033.

(5) Eason, M. G.; Pandelieva, A. T.; Mayer, M. M.; Khan, S. T.; Garcia, H. G.; Chica, R. A. Genetically Encoded Fluorescent Biosensor for Rapid Detection of Protein Expression. ACS Synth. Biol. 2020, 9 (11), 2955–2963. https://doi.org/10.1021/acssyn-bio.0c00407.

(6) Stratton, M. M.; Mitrea, D. M.; Loh, S. N. A Ca2+-Sensing Molecular Switch Based on Alternate Frame Protein Folding. ACS Chem. Biol. 2008, 3 (11), 723–732. https://doi.org/10.1021/cb800177f.

(7) Stratton, M. M.; Loh, S. N. On the Mechanism of Protein Fold-Switching by a Molecular Sensor. Proteins Struct. Funct. Bioinforma. 2010, 78 (16), 3260–3269. https://doi.org/10.1002/prot.22833.

(8) Stratton, M. M.; Loh, S. N. Converting a Protein into a Switch for Biosensing and Functional Regulation. Protein Sci. 2011, 20 (1), 19–29. https://doi.org/10.1002/pro.541.

(9) Ha, J.-H.; Shinsky, S. A.; Loh, S. N. Stepwise Conversion of a Binding Protein to a Fluorescent Switch: Application to Thermoanaerobacter Tengcongensis Ribose Binding Protein. Biochemistry 2013, 52 (4), 600–612. https://doi.org/10.1021/bi301105u.

(10) Mitrea, D. M.; Parsons, L. S.; Loh, S. N. Engineering an Artificial Zymogen by Alternate Frame Protein Folding. Proc. Natl. Acad. Sci. U. S. A. 2010, 107 (7), 2824–2829. https://doi.org/10.1073/pnas.0907668107.

(11) Do, K.; Boxer, S. G. GFP Variants with Alternative βStrands and Their Application as Light-Driven Protease Sensors: A Tale of Two Tails. J. Am. Chem. Soc. 2013, 135 (28), 10226–10229. https://doi.org/10.1021/ja4037274.

(12) Ha, J.-H.; Loh, S. N. Construction of Allosteric Protein Switches by Alternate Frame Folding and Intermolecular Fragment Exchange. Methods Mol. Biol. Clifton NJ 2017, 1596, 27–41. https://doi.org/10.1007/978-1-4939-6940-1_2.

(13) Do, K.; Boxer, S. G. Thermodynamics, Kinetics, and Photochemistry of β-Strand Association and Dissociation in a Split-GFP System. J. Am. Chem. Soc. 2011, 133 (45), 18078–18081. https://doi.org/10.1021/ja207985w.

(14) Akkapeddi, P.; Wen Teng, K.; Koide, S. Monobodies as Tool Biologics for Accelerating Target Validation and Druggable Site Discovery. RSC Med. Chem. 2021, 12 (11), 1839–1853. https://doi.org/10.1039/D1MD00188D.

(15) Sha, F.; Salzman, G.; Gupta, A.; Koide, S. Monobodies and Other Synthetic Binding Proteins for Expanding Protein Science. Protein Sci. Publ. Protein Soc. 2017, 26 (5), 910–924. https://doi.org/10.1002/pro.3148.

(16) Komuro, H.; Aminova, S.; Lauro, K.; Woldring, D.; Harada, M. Design and Evaluation of Engineered Extracellular Vesicle (EV)- Based Targeting for EGFR-Overexpressing Tumor Cells Using Monobody Display. Bioengineering 2022, 9 (2), 56. https://doi.org/10.3390/bioengineering9020056.

(17) Koide, A.; Gilbreth, R. N.; Esaki, K.; Tereshko, V.; Koide, S. High-Affinity Single-Domain Binding Proteins with a Binary-Code Interface. Proc. Natl. Acad. Sci. 2007, 104 (16), 6632–6637. https://doi.org/10.1073/pnas.0700149104.

(18) Garnish, S. E.; Meng, Y.; Koide, A.; Sandow, J. J.; Denbaum, E.; Jacobsen, A. V.; Yeung, W.; Samson, A. L.; Horne, C. R.; Fitzgibbon, C.; Young, S. N.; Smith, P. P. C.; Webb, A. I.; Petrie, E. J.; Hildebrand, J. M.; Kannan, N.; Czabotar, P. E.; Koide, S.; Murphy, J. M. Conformational Interconversion of MLKL and Disengagement from RIPK3 Precede Cell Death by Necroptosis. Nat. Commun. 2021, 12 (1), 2211. https://doi.org/10.1038/s41467-021-22400-z.

(19) Petrie, E. J.; Birkinshaw, R. W.; Koide, A.; Denbaum, E.; Hildebrand, J. M.; Garnish, S. E.; Davies, K. A.; Sandow, J. J.; Samson, L.; Gavin, X.; Fitzgibbon, C.; Young, S. N.; Hennessy, P. J.; Smith, P. P. C.; Webb, A. I.; Czabotar, P. E.; Koide, S.; Murphy, J. M. Identification of MLKL Membrane Translocation as a Checkpoint in Necroptotic Cell Death Using Monobodies. Proc. Natl. Acad. Sci. 2020, 117 (15), 8468–8475. https://doi.org/10.1073/pnas.1919960117.

(20) Miller, C. J.; McGinnis, J. E.; Martinez, M. J.; Wang, G.; Zhou, J.; Simmons, E.; Amet, T.; Abdeen, S. J.; Van Huysse, J. W.; Bowsher, R. R.; Kay, B. K. FN3-Based Monobodies Selective for the Receptor Binding Domain of the SARS-CoV-2 Spike Protein. New Biotechnol. 2021, 62, 79–85. https://doi.org/10.1016/j.nbt.2021.01.010.

(21) Kondo, T.; Iwatani, Y.; Matsuoka, K.; Fujino, T.; Umemoto, S.; Yokomaku, Y.; Ishizaki, K.; Kito, S.; Sezaki, T.; Hayashi, G.; Murakami, H. Antibody-like Proteins That Capture and Neutralize SARS-CoV-2. Sci. Adv. 2020, 6 (42), eabd3916. https://doi.org/10.1126/sciadv.abd3916.

(22) Fulcher, L. J.; Hutchinson, L. D.; Macartney, T. J.; Turnbull, C.; Sapkota, G. P. Targeting Endogenous Proteins for Degradation through the Affinity-Directed Protein Missile System. Open Biol. 2017, 7 (5), 170066. https://doi.org/10.1098/rsob.170066.

(23) Schmit, N. E.; Neopane, K.; Hantschel, O. Targeted Protein Degradation through Cytosolic Delivery of Monobody Binders Using Bacterial Toxins. ACS Chem. Biol. 2019, 14 (5), 916–924. https://doi.org/10.1021/acschembio.9b00113.

(24) Portnoff, A. D.; Stephens, E. A.; Varner, J. D.; DeLisa, M. P. Ubiquibodies, Synthetic E3 Ubiquitin Ligases Endowed with Unnatural Substrate Specificity for Targeted Protein Silencing. J. Biol. Chem. 2014, 289 (11), 7844–7855. https://doi.org/10.1074/jbc.M113.544825.

(25) Baltz, M. R.; Stephens, E. A.; DeLisa, M. P. Design and Functional Characterization of Synthetic E3 Ubiquitin Ligases for Targeted Protein Depletion. Curr. Protoc. Chem. Biol. 2018, 10 (1), 72–90. https://doi.org/10.1002/cpch.37.

(26) Carrasco-López, C.; Zhao, E. M.; Gil, A. A.; Alam, N.; Toettcher, J. E.; Avalos, J. L. Development of Light-Responsive Protein Binding in the Monobody Non-Immunoglobulin Scaffold. Nat. Commun. 2020, 11 (1), 4045. https://doi.org/10.1038/s41467-020-17837-7.

(27) He, L.; Tan, P.; Huang, Y.; Zhou, Y. Design of Smart Antibody Mimetics with Photosensitive Switches. Adv. Biol. 2021, 5 (5), e2000541. https://doi.org/10.1002/adbi.202000541.

(28) Hantschel, O. Monobodies as Possible Next-Generation Protein Therapeutics – a Perspective. Swiss Med. Wkly. 2017, No. 47. https://doi.org/10.4414/smw.2017.14545.

(29) Jin, S.; Sun, Y.; Liang, X.; Gu, X.; Ning, J.; Xu, Y.; Chen, S.; Pan, L. Emerging New Therapeutic Antibody Derivatives for Cancer Treatment. Signal Transduct. Target. Ther. 2022, 7 (1), 1–28. https://doi.org/10.1038/s41392-021-00868-x.

(30) Grebien, F.; Hantschel, O.; Wojcik, J.; Kaupe, I.; Kovacic, B.; Wyrzucki, A. M.; Gish, G. D.; Cerny-Reiterer, S.; Koide, A.; Beug, H.; Pawson, T.; Valent, P.; Koide, S.; Superti-Furga, G. Targeting the SH2-Kinase Interface in Bcr-Abl Inhibits Leukemogenesis. Cell 2011, 147 (2), https://doi.org/10.1016/j.cell.2011.08.046. 306–319.

(31) Wojcik, J.; Hantschel, O.; Grebien, F.; Kaupe, I.; Bennett, K. L.; Barkinge, J.; Jones, R. B.; Koide, A.; Superti-Furga, G.; Koide, S. A Potent and Highly Specific FN3 Monobody Inhibitor of the Abl SH2 Domain. Nat. Struct. Mol. Biol. 2010, 17 (4), 519–527. https://doi.org/10.1038/nsmb.1793.

(32) Gilbreth, R. N.; Truong, K.; Madu, I.; Koide, A.; Wojcik, J. B.; Li, N.-S.; Piccirilli, J. A.; Chen, Y.; Koide, S. Isoform-Specific Monobody Inhibitors of Small Ubiquitin-Related Modifiers Engineered Using Structure-Guided Library Design. Proc. Natl. Acad. Sci. U. S. A. 2011, 108 (19), 7751–7756. https://doi.org/10.1073/pnas.1102294108.

(33) Gupta, A.; Xu, J.; Lee, S.; Tsai, S. T.; Zhou, B.; Kurosawa, K.; Werner, M. S.; Koide, A.; Ruthenburg, A. J.; Dou, Y.; Koide, S. Facile Target Validation in an Animal Model with Intracellularly Expressed Monobodies. Nat. Chem. Biol. 2018, 14 (9), 895–900. https://doi.org/10.1038/s41589-018-0099-z.

(34) Zheng, H.; Bi, J.; Krendel, M.; Loh, S. N. Converting a Binding Protein into a Biosensing Conformational Switch Using Protein Fragment Exchange. Biochemistry 2014, 53 (34), 5505–5514. https://doi.org/10.1021/bi500758u.

(35) Limsakul, P.; Peng, Q.; Wu, Y.; Allen, M. E.; Liang, J.; Remacle, A. G.; Lopez, T.; Ge, X.; Kay, B. K.; Zhao, H.; Strongin, A. Y.; Yang, X.-L.; Lu, S.; Wang, Y. Directed Evolution to Engineer Monobody for FRET Biosensor Assembly and Imaging at Live-Cell Surface. Cell Chem. Biol. 2018, 25 (4), 370-379.e4. https://doi.org/10.1016/j.chembiol.2018.01.002.

(36) Koide, A.; Wojcik, J.; Gilbreth, R. N.; Hoey, R. J.; Koide, S. Teaching an Old Scaffold New Tricks: Monobodies Constructed Using Alternative Surfaces of the FN3 Scaffold. J. Mol. Biol. 2012, 415 (2), 393–405. https://doi.org/10.1016/j.jmb.2011.12.019.

(37) Spencer-Smith, R.; Koide, A.; Zhou, Y.; Eguchi, R. R.; Sha, F.; Gajwani, P.; Santana, D.; Gupta, A.; Jacobs, M.; Herrero-Garcia, E.; Cobbert, J.; Lavoie, H.; Smith, M.; Rajakulendran, T.; Dowdell, E.; Okur, M. N.; Dementieva, I.; Sicheri, F.; Therrien, M.; Hancock, J. F.; Ikura, M.; Koide, S.; O’Bryan, J. P. Inhibition of RAS Function through Targeting an Allosteric Regulatory Site. Nat. Chem. Biol. 2017, 13 (1), 62–68. https://doi.org/10.1038/nchembio.2231.

(38) Porebski, B. T.; Conroy, P. J.; Drinkwater, N.; Schofield, P.; Vazquez-Lombardi, R.; Hunter, M. R.; Hoke, D. E.; Christ, D.; McGowan, S.; Buckle, A. M. Circumventing the Stability-Function Trade-off in an Engineered FN3 Domain. Protein Eng. Des. Sel. PEDS 2016, 29 (11), 541–550. https://doi.org/10.1093/pro-tein/gzw046.

(39) Batori, V.; Koide, A.; Koide, S. Exploring the Potential of the Monobody Scaffold: Effects of Loop Elongation on the Stability of a Fibronectin Type III Domain. Protein Eng. Des. Sel. 2002, 15 (12), 1015–1020. https://doi.org/10.1093/protein/15.12.1015.

(40) Radley, T. L.; Markowska, A. I.; Bettinger, B. T.; Ha, J.-H.; Loh, S. N. Allosteric Switching by Mutually Exclusive Folding of Protein Domains. J. Mol. Biol. 2003, 332 (3), 529–536. https://doi.org/10.1016/s0022-2836(03)00925-2.

(41) Cutler, T. A.; Mills, B. M.; Lubin, D. J.; Chong, L. T.; Loh, S. N. Effect of Interdomain Linker Length on an Antagonistic Folding-Unfolding Equilibrium between Two Protein Domains. J. Mol. Biol. 2009, 386 (3), 854–868. https://doi.org/10.1016/j.jmb.2008.10.090.

(42) Nguyen, A. W.; Daugherty, P. S. Evolutionary Optimization of Fluorescent Proteins for Intracellular FRET. Nat. Biotechnol. 2005, 23 (3), 355–360. https://doi.org/10.1038/nbt1066.

(43) Shaner, N. C.; Lambert, G. G.; Chammas, A.; Ni, Y.; Cranfill, P. J.; Baird, M. A.; Sell, B. R.; Allen, J. R.; Day, R. N.; Israelsson, M.; Davidson, M. W.; Wang, J. A Bright Monomeric Green Fluorescent Protein Derived from Branchiostoma Lanceolatum. Nat. Methods 2013, 10 (5), 407–409. https://doi.org/10.1038/nmeth.2413.

(44) Nagai, T.; Ibata, K.; Park, E. S.; Kubota, M.; Mikoshiba, K.; Miyawaki, A. A Variant of Yellow Fluorescent Protein with Fast and Efficient Maturation for Cell-Biological Applications. Nat. Biotechnol. 2002, 20 (1), 87–90. https://doi.org/10.1038/nbt0102-87.

(45) John, A. M.; Sekhon, H.; Ha, J.-H.; Loh, S. N. Engineering a Fluorescent Protein Color Switch Using Entropy-Driven β-Strand Exchange. ACS Sens. 2022, 7 (1), 263–271. https://doi.org/10.1021/acssensors.1c02239.

(46) Gil, A. A.; Carrasco-López, C.; Zhu, L.; Zhao, E. M.; Ravindran, P. T.; Wilson, M. Z.; Goglia, A. G.; Avalos, J. L.; Toettcher, J. E. Optogenetic Control of Protein Binding Using Light-Switchable Nanobodies. Nat. Commun. 2020, 11 (1), 4044. https://doi.org/10.1038/s41467-020-17836-8.

(47) Nicholes, N.; Date, A.; Beaujean, P.; Hauk, P.; Kanwar, M.; Ostermeier, M. Modular Protein Switches Derived from Antibody Mimetic Proteins. Protein Eng. Des. Sel. 2016, 29 (2), 77–85. https://doi.org/10.1093/protein/gzv062.

(48) Forrer, P.; Stumpp, M. T.; Binz, H. K.; Plückthun, A. A Novel Strategy to Design Binding Molecules Harnessing the Modular Nature of Repeat Proteins. FEBS Lett. 2003, 539 (1–3), 2–6. https://doi.org/10.1016/s0014-5793(03)00177-7.

(49) Stumpp, M. T.; Forrer, P.; Binz, H. K.; Plückthun, A. Designing Repeat Proteins: Modular Leucine-Rich Repeat Protein Libraries Based on the Mammalian Ribonuclease Inhibitor Family. J. Mol. Biol. 2003, 332 (2), 471–487. https://doi.org/10.1016/s0022-2836(03)00897-0.

(50) Quijano-Rubio, A.; Yeh, H.-W.; Park, J.; Lee, H.; Langan, R. A.; Boyken, S. E.; Lajoie, M. J.; Cao, L.; Chow, C. M.; Miranda, M. C.; Wi, J.; Hong, H. J.; Stewart, L.; Oh, B.-H.; Baker, D. De Novo Design of Modular and Tunable Protein Biosensors. Nature 2021, 591 (7850), 482–487. https://doi.org/10.1038/s41586-021-03258-z.

(51) Langan, R. A.; Boyken, S. E.; Ng, A. H.; Samson, J. A.; Dods, G.; Westbrook, A. M.; Nguyen, T. H.; Lajoie, M. J.; Chen, Z.; Berger, S.; Mulligan, V. K.; Dueber, J. E.; Novak, W. R. P.; El-Samad, H.; Baker, D. De Novo Design of Bioactive Protein Switches. Nature 2019, 572 (7768), 205–210. https://doi.org/10.1038/s41586-019-1432-8.

(52) Dixon, A. S.; Schwinn, M. K.; Hall, M. P.; Zimmerman, K.; Otto, P.; Lubben, T. H.; Butler, B. L.; Binkowski, B. F.; Machleidt, T.; Kirkland, T. A.; Wood, M. G.; Eggers, C. T.; Encell, L. P.; Wood, K. V. NanoLuc Complementation Reporter Optimized for Accurate Measurement of Protein Interactions in Cells. ACS Chem. Biol. 2016, 11 (2), 400–408. https://doi.org/10.1021/acschembio.5b00753.

(53) Chandler, P. G.; Tan, L. L.; Porebski, B. T.; Green, J. S.; Riley, B. T.; Broendum, S. S.; Hoke, D. E.; Falconer, R. J.; Munro, T. P.; Buckle, M.; Jackson, C. J.; Buckle, A. M. Mutational and Biophysical Robustness in a Prestabilized Monobody. J. Biol. Chem. 2021, 296. https://doi.org/10.1016/j.jbc.2021.100447.

(54) Pichler, A.; Gast, A.; Seeler, J. S.; Dejean, A.; Melchior, F. The Nucleoporin RanBP2 Has SUMO1 E3 Ligase Activity. Cell 2002, 108 (1), 109–120. https://doi.org/10.1016/S0092-8674(01)00633-X.

(55) Shinsky, S. A.; Hu, M.; Vought, V. E.; Ng, S. B.; Bamshad, M. J.; Shendure, J.; Cosgrove, M. S. A Non-Active Site SET Domain Surface Crucial for the Interaction of MLL1 and the RbBP5-ASH2L Heterodimer within MLL Family Core Complexes. J. Mol. Biol. 2014, 426 (12), 2283–2299. https://doi.org/10.1016/j.jmb.2014.03.011.

(56) Patel, A.; Dharmarajan, V.; Vought, V. E.; Cosgrove, M. S. On the Mechanism of Multiple Lysine Methylation by the Human Mixed Lineage Leukemia Protein-1 (MLL1) Core Complex. J. Biol. Chem. 2009, 284 (36), 24242–24256. https://doi.org/10.1074/jbc.M109.014498.

(57) Pomorski, A.; Kochańczyk, T.; Miłoch, A.; Krężel, A. Method for Accurate Determination of Dissociation Constants of Optical Ratiometric Systems: Chemical Probes, Genetically Encoded Sensors, and Interacting Molecules. Anal. Chem. 2013, 85 (23), 11479–11486. https://doi.org/10.1021/ac402637h.

