## Supplemental Information for "An adaptable, monobody-based biosensor scaffold with FRET output"

**TABLE OF CONTENTS**

**Figure S1:** **Amino acid sequences of FN3-AFF switches.
Figure S2: Stability tuning of N-fold and cp-fold.
Figure S3: Binding affinities of FN3 and cpFN3 and validation of binding-null mutations.
Figure S4: Characterization of FN3-AFF switches by size exclusion chromatography.
Figure S5: FN3-AFF^Ras^ bound h-Ras with K_d_ = 423 ± 200 nM.
Figure S6: FN3^WDR5^ retained ligand binding activity after insertion of cpYFP.
Figure S7: The turn-on rate of FN3-AFF^GE^ did not depend on WDR5 concentration.**

**A**

```
SH2  MCGEWKEVTVPYSSSAYTVTGLKPGTEYEFRVYAWGEDSAGAAFMYSPSSVSVTTGGSGS 60
SUMO MCGEWKEVTVPYSSSAYTVTGLKPGTEYEFRVYAA-----YASTWYSPSSVSVTTGGSGS 55
WDR5 MCGEWQKVKVPYSSSAYTVTGLKPGTEYEFRVYAYQGGGAWHP-YGSPSSVSVTTGGSGS 59
hRas MCGEWQKVEVPYSSSAYTVTGLKPGTEYEFRVYAWGWHGQVYA-AMSPSSVSVTTGGSGS 57
  *****::* *************************            **************
DE loop                    FG loop
SH2  GGASGGATGGSGGGSPSVPGNLRVTDVTSTSVTLSWDAPMSSSSVYYYRVEYREAGCGEW 120
SUMO GGASGGATGGSGGGSPSVPGNLRVTDVTSTSVTLSWDAGR--WFVEYYRVEYREAGCGEW 113
WDR5 GGASGGATGGSGGGSPSVPGNLRVTDVTSTSVTLSWDAPAV--TVVHYRVEYREAGCGEW 117
hRas GGASGGATGGSGGGSPSVPGNLRVTDVTSTSVTLSWDAPAV--TVDYYRVEYREAGCGEW 115
 **************************************      * :*************
                         BC loop
SH2  KEVTVPYSSSAYTVTGLKPGTEYEGRVYAWGEDSAGYMFMYSPSSVSVTTLEHHHHHHHH 180
SUMO KEVTVPYSSSAYTVTGLKPGTEYEGRVYAHY-----WSTWYSPSSVSVTTLEHHHHHHHH 168
WDR5 QKVKVPYSSSAYTVTGLKPGTEYEGRVYAYQGGGR-WHPYGSPSSVSVTTLEHHHHHHHH 176
hRas QKVEVPYSSSAYTVTGLKPGTEYEGRVYAWGWHGQVYY-YMSPSSVSVTTLEHHHHHHHH 174
 ::* *************************       :    *******************
          DE loop                    FG loop
```

**B**

**MGSKGEELFGGIVPILVELEGDVNGHKFSVSGEGEGDATYGKLTLKFICT 50
TGKLPVPWPTLVTTLTWGVQCFSRYPDHMKQHDFFKSVMPEGYVQERTIF 100
FKDDGNYKTRAEVKFEGDTLVNRIELKGIDFKEDGNILGHKLEYNYISHN 150
VYITADKQKNGIKANFKARHNITDGSVQLADHYQQNTPIGDGPVILPDNH 200
YLSTQSALSKDPNEKRDHMVLLEFVTAAGITHGMDELYKGGGSGGEWQKV 250
KVPYSSSAYTVTGLKPGTEYEFRVYAYQGGGAWHPYGSPSSVSVTTGGSG 300
SGGASGGATGGSGGGSPSVPGNLRVTDVTSTSVTLSWDAPAVTVVHYRVE 350
YREAGLPDNHYLSYQSVLSKDPNEKRDHMVLLEFVTAAGITLGMDELYKG 400
GGSGGMVSKGEELFTGVVPILVELDGDVNGHKFSVRGEGEGDATNGKLTL 450
KLICTTGKLPVPWPTLVTTLGYGLACFSRYPDHMKQHDFFKSAMPEGYVQ 500
ERTISFKDDGTYKTRAEVKFEGDTLVNRIELKGIDFKEDGNILGHKLEYN 550
FNSHNVYITADKQKNGIKANFKIRHNVEDGSVQLADHYQQNTPIGDGPVL 600
GEWQKVKVPYSSSAYTVTGLKPGTEYEGRVYAYQGGGRWHPYGSPSSVSV 650
TTLEHHHHHHHH**

**Figure S1:** **Amino acid sequences of FN3-AFF switches.** **(A)** FN3-AFF^SH2^, FN3-AFF^SUMO^, and FN3-AFF^WDR5^ are shown with colors corresponding to regions shown in Figure 1, as follows: red, duplicated sequence (cp-fold); black, 18-AA linker; purple, shared sequence; blue, duplicated sequence (N-fold); grey, HisTag purification sequence. CDR-like loops are indicated in bars below the sequences. Underlined residues in the FG loop (here shown in the cp-fold; the analogous residues were mutated in the N-fold to generate FN3-AFF^rev^ switches) denote binding-null mutations. The green asterisk indicates the Phe to Gly tuning mutation. **(B)** The sequence of FN3-AFF^GE^ is shown with the same color coding as in panel A, and with CyPet in green and cpYFP in yellow, and linkers underlined in black.

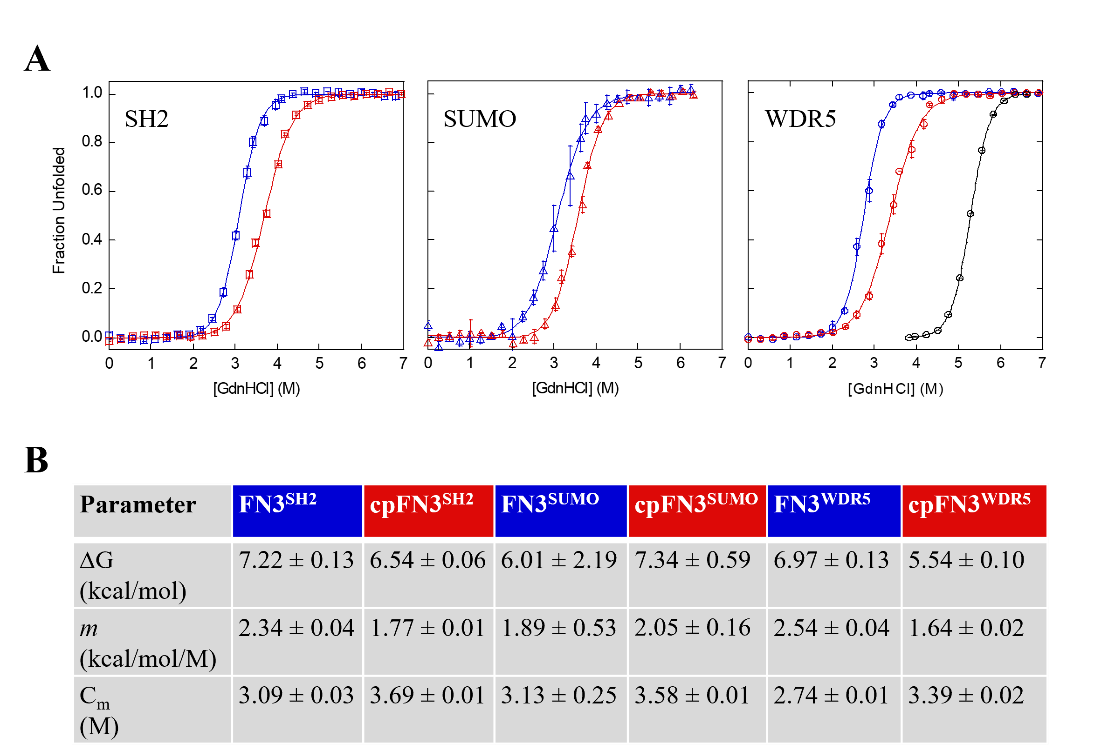

**Figure S2: Stability tuning of N-fold and cp-fold. (A)** GdnHCl denaturation curves of FN3 (blue) and cpFN3^bn^ (red) are shown, with the ligands to which they bind indicated in the insets. FN3 proteins contain the tuning mutation shown in Figure S1. Black data represent FN3^WDR5^ prior to introducing the tuning mutation. Trp emission spectra (300 – 450 nm with excitation at 280 nm) were recorded on a Horiba SpectraMax 4 spectrofluorimeter and fit using Igor Pro (Wavemetrics, Inc.) to obtain the peak center. This value was plotted as a function of [GdnHCl] and fit to the linear extrapolation equation [Santoro M.M. and Bolen D.W. (1988), *Biochemistry 27*, 8063-8068] to obtain thermodynamic parameters shown in **(B)**. Conditions were 2 μM protein, 20 mM Tris (pH 7.4), 0.3 M NaCl, 0.005 % TWEEN-20, 22 °C. Error bars in panel A and uncertainty values in panel B are s.d. (n = 3).

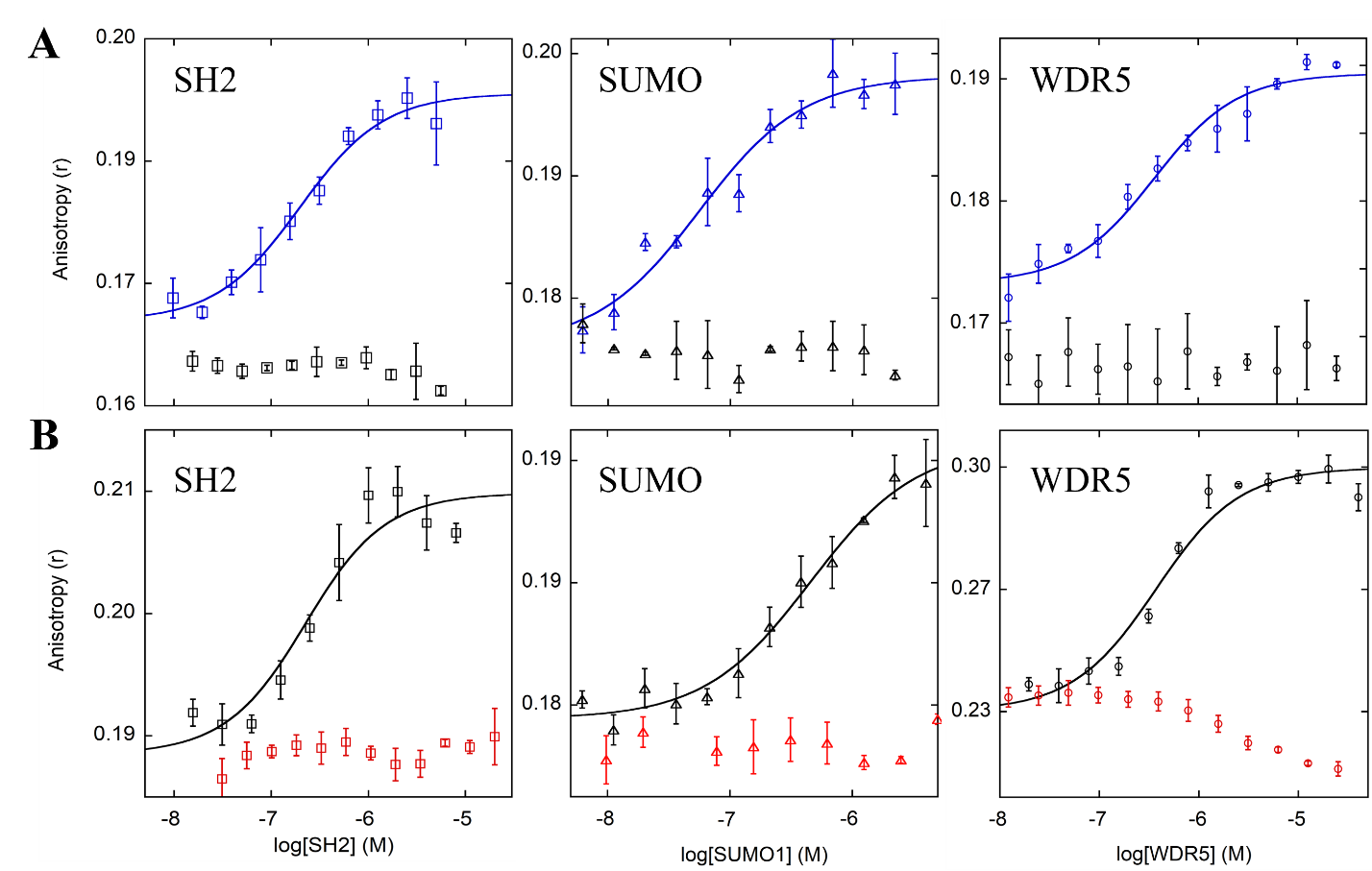

**Figure S3: Binding affinities of FN3 and cpFN3 and validation of binding-null mutations.** **(A)** Binding of FN3 (blue) and FN3^bn^ (black) to their respective ligands (insets) was determined by fluorescence anisotropy. **(B)** Binding of cpFN3 (black) and cpFN3^bn^ (red) to their respective ligands (insets) was monitored by fluorescence anisotropy. In panels A and B, the FN3 (blue) and cpFN3^bn^ (red) components make up the FN3-AFF switch shown in Fig. 1B. For all data sets except cpFN3^SH2^, FN3 and cpFN3 were labeled at the Cys positions in the main text using fluorescein 5 maleimide, and 0.25 μM labeled protein was mixed with unlabeled ligands at the indicated concentrations. Samples were equilibrated for 15 min at room temperature and then fluorescence anisotropy was measured on a SpectraMax plate reader as described in the Methods. The cpFN3^SH2^ data set was obtained by labeling SH2 at its N-terminus (using fluorescein NHS ester, ThermoFisher Scientific) and titrating in unlabeled cpFN3^SH2^. Buffer was 20 mM Tris (pH 7.4), 0.3 M NaCl, 0.005% Tween-20, and 1 mM TCEP. Anisotropy data were fit to the quadratic binding equation (solid lines). Error bars are s.d. (n = 3).

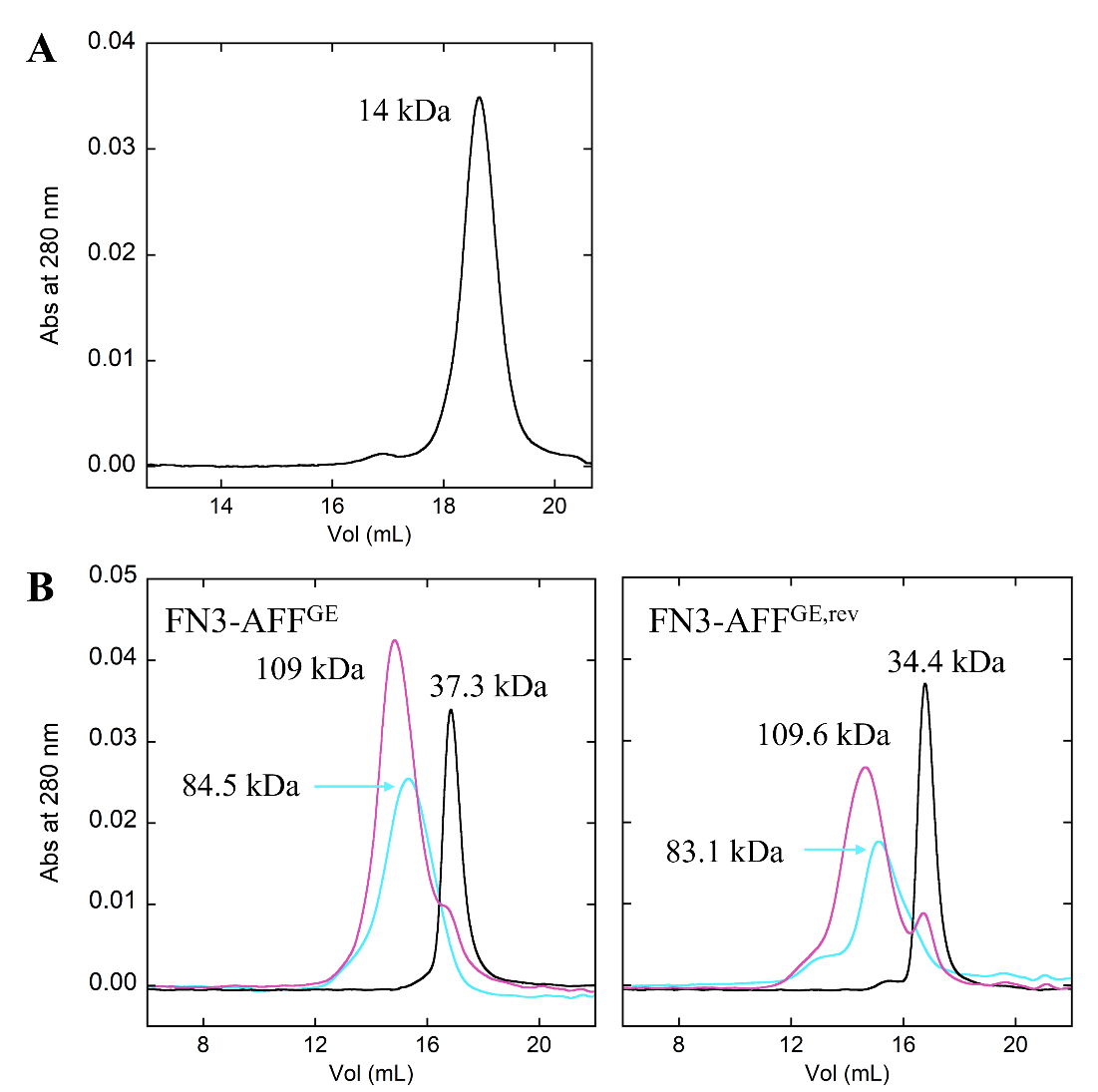

**Figure S4: Characterization of FN3-AFF switches by size exclusion chromatography. (A)** FN3-AFF^WDR5^ was >95 % pure and monomeric. **(B)** FN3-AFF^GE^ was >95% pure and monomeric, and binds WDR5. FN3-AFF^GE^ is shown in cyan, WDR5 in black, and the mixture of FN3-AFF^GE^ and WDR5 in purple. All protein concentrations were 20 μM. Buffer conditions were 20 mM Tris (pH 7.4), 0.3 M NaCl, 0.005% Tween-20, and 1 mM TCEP. Binding samples were incubated for 10 min at room temperature and then injected onto a Superdex 200 Increase 10/300GL column (Cytiva, Marlborough MA) using a Bio-Rad DuoFlow chromatography system.

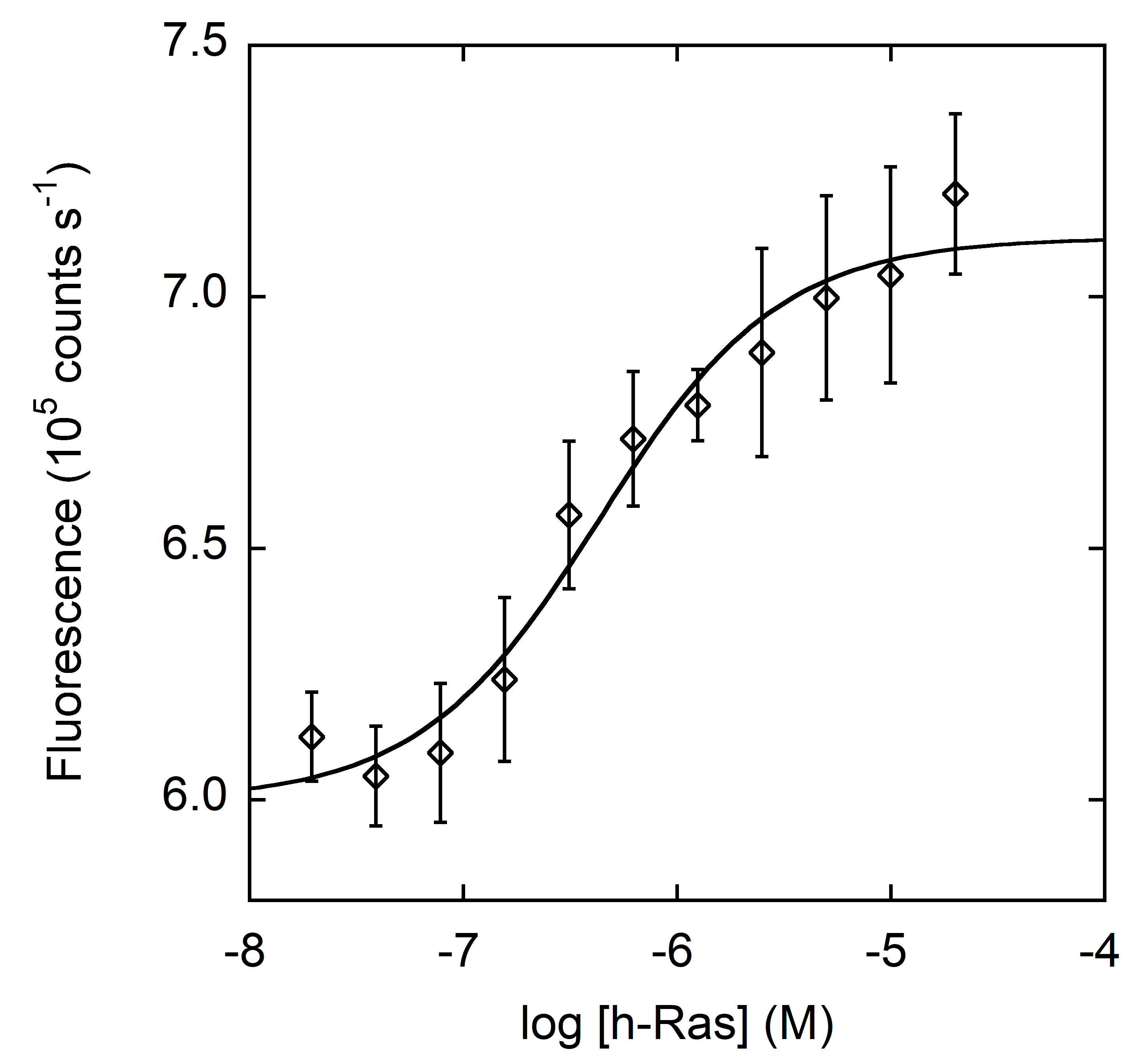

**Figure S5: FN3-AFF^Ras^ bound h-Ras with K_d_ = 423 ± 200 nM.** BODIPY-labeled FN3-AFF^Ras^ (50 nM) was mixed with h-Ras and equilibrated for 12 h at 22 °C before being scanned on a Fluoromax-4 spectrofluorometer as described in the methods. Fluorescence data (510 nm) were fit to the quadratic binding equation (solid line). Error bars are s.d. (n = 3).

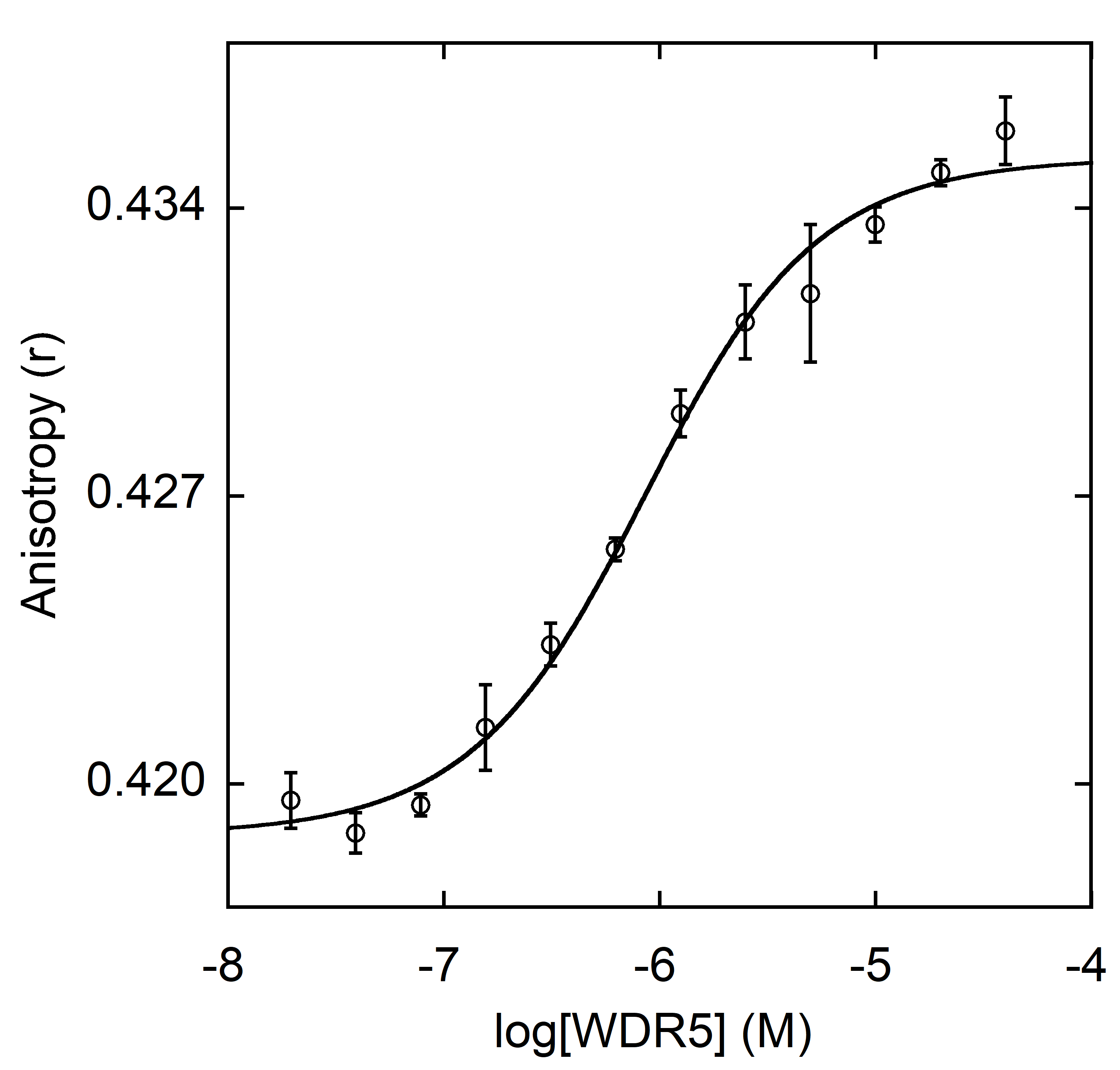

**Figure S6: FN3^WDR5^ retained ligand binding activity after insertion of cpYFP.** The cpYFP-FN3^WDR5^ fusion protein was mixed with WDR5 and allowed to equilibrate for 15 min at room temperature, whereupon fluorescence anisotropy was measured on a SpectraMax plate reader as described in the methods. Fitting the data to the quadratic binding equation (line) yielded K_d_ = 727.1 ± 52.9 nM. This value is ~2-fold higher than that of FN3^WDR5^ (Figure S3A).

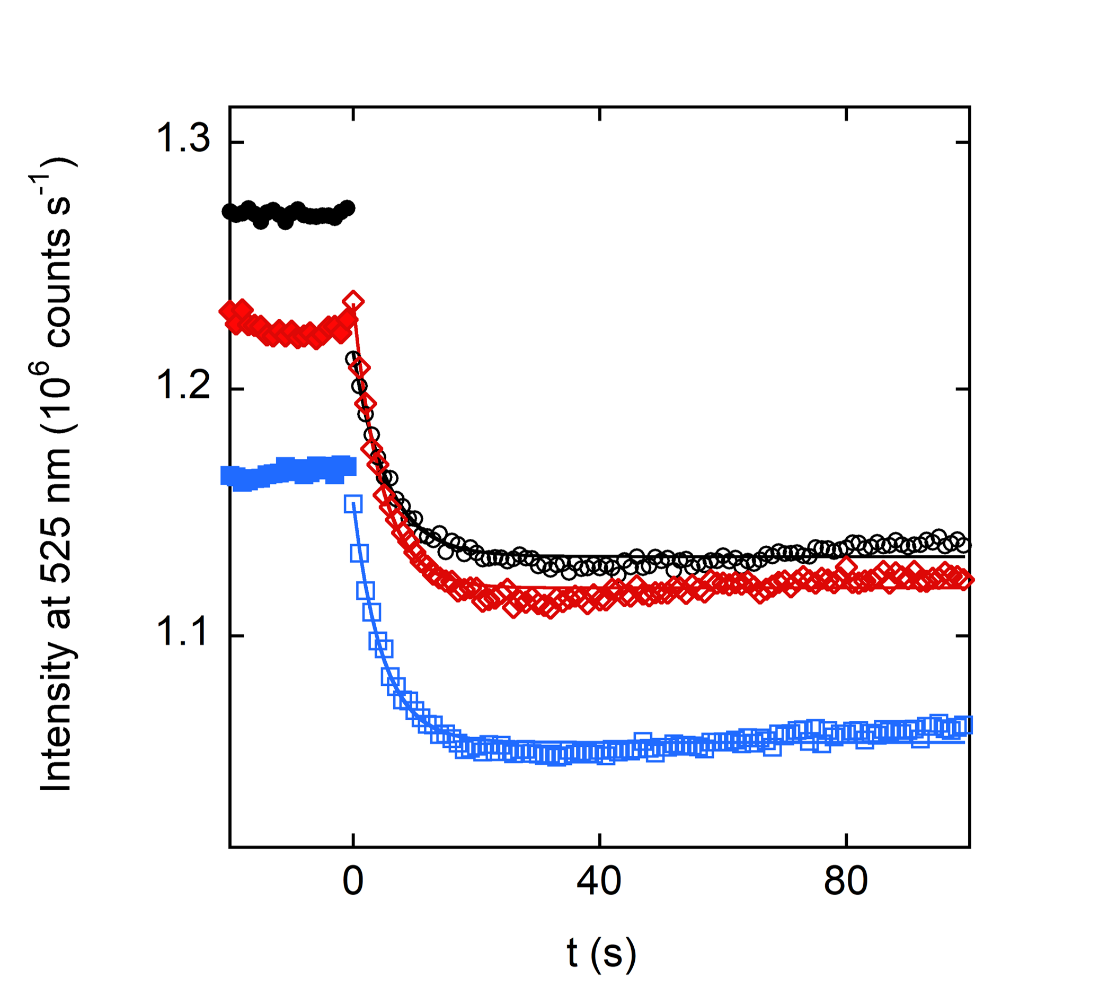

**Figure S7: The turn-on rate of FN3-AFF^GE^ did not depend on WDR5 concentration.** After recording baseline fluorescence of FN3-AFF^GE^ (0.5 μM) for ~30 s (closed symbols), WDR5 was added at time zero to 5 μM (open red diamonds), 10 μM (open black circles), or 20 μM (open blue squares). Rates were obtained by fitting the decrease in cpYFP emission to single exponential functions (solid lines). Conditions were 20 mM Tris (pH 7.4), 0.3 M NaCl, 0.005% Tween-20, and 1 mM TCEP (22 °C).
